# An ‘Epidemic Diversity’ conceptual model explains how host genetic diversity affects variation in parasite success

**DOI:** 10.1101/2024.05.28.596150

**Authors:** Sam Paplauskas

**Author notes:** Corresponding author: (SP).

## Abstract

According to conventional wisdom, disease transmission rate is usually higher in more genetically homogenous host populations. Previous studies have principally considered how host population genetic diversity impacts the mean of parasite infection performance. However, when considering risks from epidemics and the emergence of novel infectious diseases, variability in parasite success may be just as important as the mean. Here we propose an Epidemic Diversity Model for how host-parasite population genetic diversity influences both the mean and variability in parasite success. We evaluate this conceptual model by re-analysing effect size data from two meta-analyses, including 211 comparisons of high versus low genetic diversity host populations from 48 studies. Our analysis challenges previous understanding by demonstrating that high host population genetic diversity only reduces mean parasite success for specialist parasites with narrow host range, but not for generalist multi-host parasites. We also find that the combination of host range and parasite population genetic diversity determine the effect of host population genetic diversity on the variability in parasite success. These results have important implications for the management of host population genetic diversity for natural populations.

## Introduction

Epidemics can be defined as extremely rapid increases in disease prevalence over the endemic (baseline) level of disease [1]. They pose a major threat to food security, biodiversity and human health [2]. This is because epidemics can devastate crop harvests [3], threaten local host populations with disease-induced extinction [4] and increase the risk of zoonotic spillover [5]. Previous research suggests that less genetically diverse host populations tend to experience larger epidemics [6]. Although this phenomenon is well-established in the agricultural literature and is often referred to as a ‘monoculture effect’ [3,7,8], it is much less well-understood how the level of genetic diversity in animal and bacterial species (i.e. not-plant hosts) determines epidemic size in nature.

Previous studies by Ekroth et al. [9] and Gibson and Nguyen [10] found that higher levels of host genetic diversity reduced the mean for various metrics of parasite success (including disease prevalence, parasite load and virulence) across a range of both animal and bacterial host-parasite systems. This evidence is centred around means, and although disease prevalence is only one measure of parasite success, it appears to support the conventional wisdom that less genetically diverse host populations tend to experience larger epidemics, at least *on average* [6]. However, despite this important discovery [9,10], many host populations in nature experience epidemics that *vary* dramatically in both time and space [11–18]. Therefore, understanding not only the mean, but also the *variability* of parasite success is important.

Previous research suggests that the understanding the variability of parasite success may be important from the perspective that high levels of variation are akin to particularly large, or severe epidemics. For example, a highly cited paper published by Tarpy [19] in *Nature* showed that more genetically diverse honeybee colonies had less variable disease prevalence. As such, they suggested that this could potentially protect more diverse honeybee colonies against the occurrence of more severe epidemics. Alternatively, increased understanding of the variability of parasite success could increase our general understanding of the repeatability of disease experiments, which are characterised by large amounts of variation [20].

Any relationship between host population genetic diversity and either the mean or variability in parasite success will likely depend on the corresponding level of parasite population genetic diversity and infection genetics. For example, assuming that there is some level of specificity for infection [21], where host and parasite genotypes vary in their susceptibility or infectivity to one another, individual host genotypes are more likely to be infected by a diverse parasite population. Therefore, the parasite transmission rate and the overall disease prevalence should be higher for more diverse parasite populations [22,23]. However, the amount of host-parasite specificity for infection varies between different models of infection genetics [24]. In these models, infection specificity ranges from matching allele paradigms where one parasite genotype can only infect a single host genotype, to gene-for-gene interactions where infectious parasite genotypes are able to infect many or all host genotypes [24]. In reality, these models represent opposite ends of a continuum and host-parasites systems vary in where they fall along this continuum [24]. Nevertheless, the strength of specificity clearly has the potential to modulate how host-parasite population genetic diversity influences both mean and variation in parasite infection success.

Here, we propose an ‘Epidemic Diversity’ conceptual model to describe how host population genetic diversity and parasite population genetic diversity may influence both the mean and variability of parasite success (Fig 1). We envisage a metapopulation structure composed of multiple local populations. Local populations vary in their level of genetic diversity (measured at the inter-, versus intra-, group level; Fig S1). Low genetic diversity results in local populations that harbour few host or parasite genotypes, but the identity of these genotypes may differ between local populations. High genetic diversity increases the number of genotypes present and local populations are less genetically differentiated. Each host individual encounters many potentially infectious parasites. Initially we consider a case of high infection specificity, where close matching is required between host and parasite genotypes for successful infection. We predict the following patterns in mean and variance of parasite success across the local populations:

A) When host and parasite population genetic diversity are both low (Fig 1A), parasite success will be highly variable because host populations will be composed either of mostly susceptible or mostly resistant host genotypes. Mean parasite success will be intermediate, and determined by the relative frequency of resistant and susceptible populations.
B) When genetic diversity is high in the host population but low in the parasite population (Fig 1B) both mean and variability in parasite success will be low because the parasite’s encounter rate with matching host genotypes will be consistently low.
C) When genetic diversity is low in the host population but high in the parasite population (Fig 1C) mean parasite success will be high and variability in parasite success low, because many hosts will encounter matching parasite genotypes and efficient chains of transmission will establish.
D) When genetic diversity is high in both host and parasite populations (Fig 1D) the mean and variability in parasite success will be intermediate because although the probability of hosts encountering infectious parasite genotypes is high, onward transmission is impaired by the diversity of host genotypes present.

**Fig 1.**
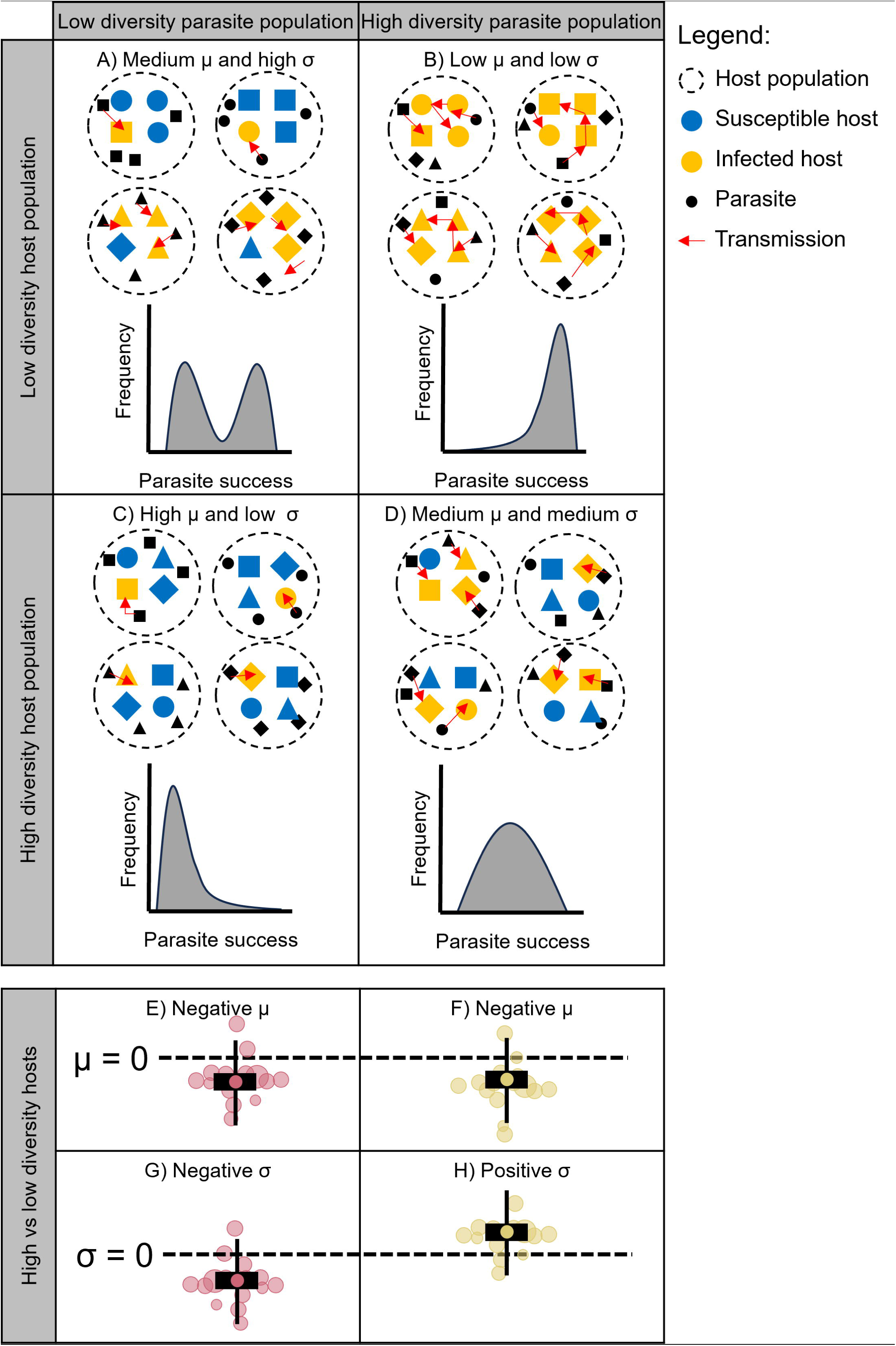
A hypothetical ‘Epidemic Diversity’ model for the combined relationship between host and parasite population genetic diversity and either the mean (μ) or the variability (. O’ **) in parasite success.** There are four hypothetical populations for each combination of host and parasite population genetic diversity (dashed circles). The level of population genetic diversity is indicated by the number of unique host and parasite genotypes (large and small shapes respectively) and is the same in each replicate population. The colour of hosts indicates their infection status, such that susceptible hosts are white and infected hosts are black, whereas the parasite is always the same colour (also black). Parasite transmission can only occur between matching host and parasite genotypes (shapes) and is indicated by the red arrows. The resulting frequency distributions of parasite infection success for each set of replicate populations is shown at the bottom of each plot. Notably, this hypothetical model only applies for host--parasite systems that have a high level of genetic specificity for infection (i.e. matching-allele versus gene-for-gene infection genetics [24]). The bottom panel of the facetted grid show the expected result from four comparisons of high versus low genetic diversity host populations (two x low parasite diversity, two x high parasite diversity). The design of these plots mirrors the orchard plots for the meta-analytical models from Fig 5.

Our model predicts that under this scenario of high genetic specificity for successful infection, an increase in host population genetic diversity will result in a decline in *mean* parasite infection success regardless of the parasite population’s genetic diversity (Fig 1E and F). However, increased host population genetic variation will cause reduced *variability* in parasite infection success when parasite genetic diversity is low (Fig 1G) but increased *variability* when parasite diversity is high (Fig 1F). All these effects will be sensitive to the genetic specificity that underpins the infection process. In cases where parasite virulence genotypes can infect many (or all) host genotypes (sensu gene-for-gene models) these predictions should break down and the effects of host population genetic diversity on parasite infection success should disappear.

This Epidemic Diversity conceptual model builds on previous work by Bensch et al. [25], who considered three out of four of the combinations in Fig 1. We extend this previous work by considering how high parasite genetic diversity can lead to more consistent parasite success in genetically depauperate host populations (Fig 1C). Furthermore, in our work we consider how these relationships between host-parasite population genetic diversity and parasite success depend on the strength of genetic specificity for infection success.

It is impossible to make an explicit test of the effect of genetic specificity for infection success in our analysis because genetic specificity is not characterised in most host-parasite systems (but see [26]). However, we investigate the impact of a proxy for genetic infection susceptibility: parasite host-range. This is because it is only specialist parasites (with a narrow host range) that are likely to have undergone tight coevolution with their host species of the type that promotes the maintenance of matching-allele-type genetic variants under negative frequency dependent selection. Whereas for generalist parasites that infect many host species (and have not tightly coevolved with any single host), virulence mechanisms are more likely to be general and confer ability to infect many host genotypes.

To evaluate our proposed Epidemic Diversity conceptual model, we combine the data from two previous meta-analyses that investigated the effect of host population genetic diversity on mean metrics of parasite infection success [9,10]. By pooling the data from these original meta-analyses, we achieve a bigger sample size so that we can investigate patterns in parasite infection success variance, as well as the impact of additional moderator variables. Since we focussed on the re-analysis of previous studies by Ekroth et al. [9] and Gibson and Nguyen [10], which mainly focussed on the effect of differences in host genetic diversity in animal and bacterial populations, rather than plant populations, we also focussed our study on animal and bacterial populations. If a researcher was interested in investigating the effect of differences in host genetic diversity in plant populations (*sensu* [27]), a much more comprehensive review of the literature would be required.

## Materials and Methods

### Summary

We combined the data from two previous meta-analyses [9,10] that used the standardised mean difference (SMD) to calculate the significance of the relationship between host genetic diversity and mean metrics of parasite infection success. These data were mainly from studies where the metrics of parasite infection success were measured from replicate host populations that were already classified qualitatively as having either a ‘high’ or ‘low’ level of genetic diversity (27 out a total of 48 independent studies), including studies of replicate host populations with a high versus low level of inbreeding (e.g. [28]), different combinations of host genotypes (e.g. [29]) or comparisons between wild host populations exposed to different selection regimes (e.g. a population bottleneck, random genetic drift, e.g. [30]). However, there were some studies that assessed a continuous measure of host population genetic diversity (for the absence of any significant correlation between mean metrics of parasite infection success and host population genetic diversity, involving all of the studies from this subset of data, see [10]), and therefore these data were binned into ‘high’ and ‘low’ categories. Following Ekroth et al. [9] and Gibson & Nguyen [10] we used the SMD to assess factors influencing mean parasite infection success; we then extended these previous analyses by using the log coefficient of variation ratio (lnCVR) to assess factors influencing variation in parasite success. In the original studies, this variation was quantified in various ways: between experimental replicates in a laboratory environment; between multiple natural host populations with similar genetic diversity; or sometimes between repeated measures of single populations along a time series.

### Data collection

Data collection for each comparison of a group of high versus low genetic diversity host populations involved five steps (Fig 2):

1) *Combination*: First, we combined the list of studies from [9] and [10], removed any duplicate studies and added the data used to calculate the effect size on mean parasite success (SMD) and its sampling variance in the original studies (mean, standard deviation, sample size), the metric of parasite success and the unique study, experiment and replicate identifiers used to account for the non-independence of separate effect sizes. We did not use the parasite success data from Ekroth et al. [9] because the original data extracted from each study was not included in their online supplementary material and this meant we would have been unable to check the data accuracy during the third step of data collection (validation). However, we extracted any missing data from these studies and recalculated the mean, standard deviation and sample size ourselves during the fourth and fifth steps of data collection (extraction and recalculation).

**Fig 2.**
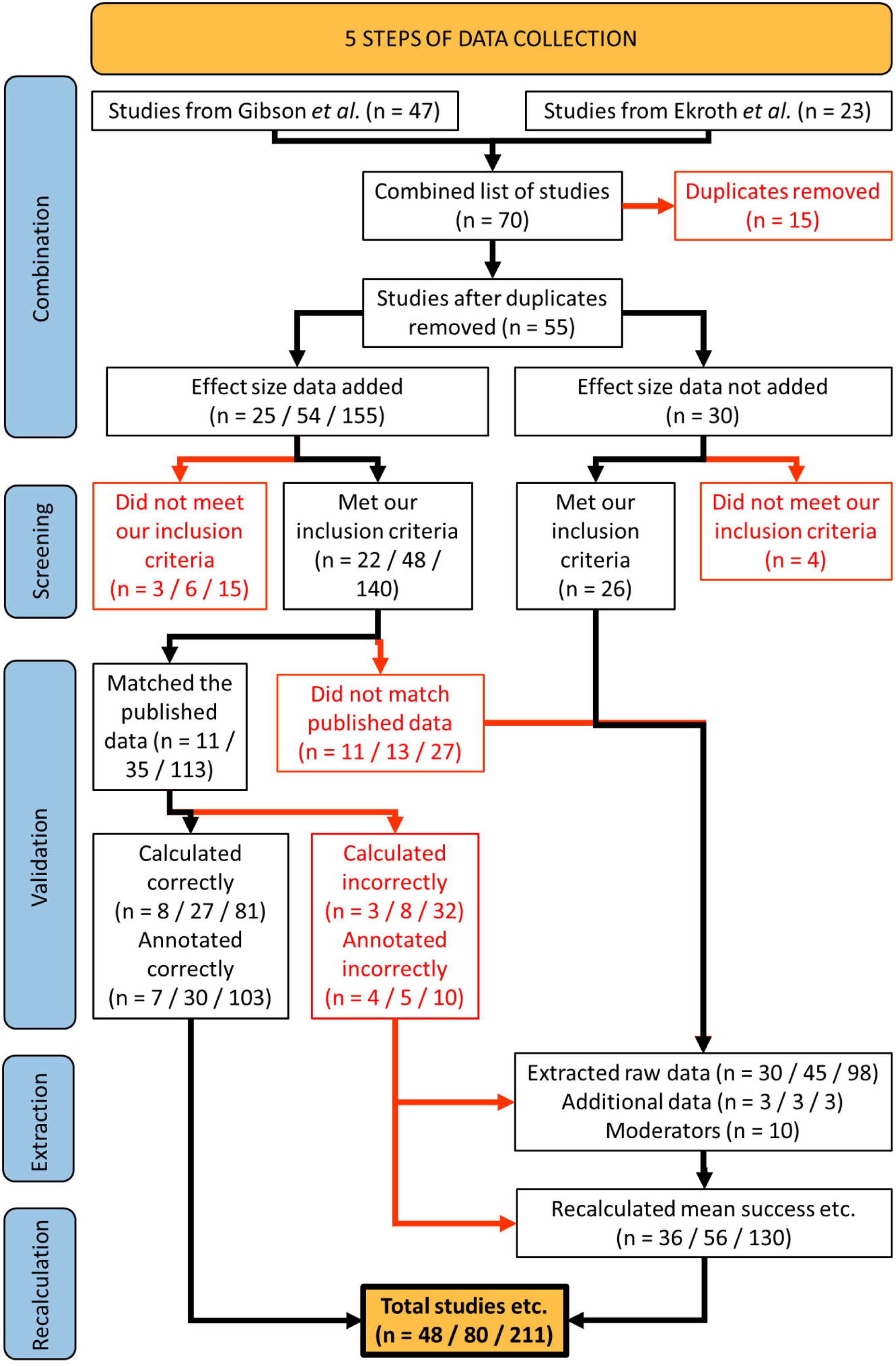
How data was collected for each comparison of a parasite infection success metric between high versus low genetic diversity groups of host populations, which was later used in calculating effect sizes. n = number of studies / experiments / comparisons (the multi-level structure of the data make it appear that some sums are incorrect). Adapted from the preferred reporting items for systematic reviews and meta-analyses (PRIMSA) statement [55].

The data used to calculate Fisher’s z (an effect size for the difference between two correlation coefficients) for the observational field studies from Gibson & Nguyen [10], was not available in the correct format to calculate either SMD or lnCVR. Therefore, we did not include this information (from multiple populations with a continuous measure of genetic diversity) at this stage and instead extracted the data from the original publications ourselves and recalculated it during steps 4 and 5. Also, we excluded any studies on plants (wild or agricultural) because a more detailed analysis of the plant literature would require a separate review.

2) *Screening*: Second, we amended the inclusion criteria used in the original meta-analyses [9,10] (Table S1) and removed any studies, experiments or comparisons which did not meet these criteria:

(i) ‘Parasite success’, which we define as the ability of a parasite to spread among hosts (transmission rate, infection rate, prevalence), replicate on / within hosts (macro / microparasite load, disease severity) or kill hosts (virulence i.e. host survival / mortality rate) was measured among replicate populations across time or space.
(ii) Parasite success data was collected from two or more host populations with a difference in genetic diversity assessed by metrics such as: individual inbreeding status (inbred versus outbred), genotypic diversity (high versus low) or heterozygosity (high vs low).
(iii) Genetic diversity was quantified at the level of the host population, rather than for community-level diversity.
(iv) The study focused on an animal or bacterial host species.
(v) The study did not re-analyse the data from a previously published study.
(vi) The parasite success data was not replicated simply by using an alternate way of measuring host population diversity.
(vii) Figures required to extract parasite success data were clearly legible.

(3) *Validation*: Third, we checked the accuracy of the data from the excel spreadsheets used to calculate the summary of the parasite data for each group of host populations with either high or low genetic diversity from the online data supplied by one of the previous meta-analyses [10] and corrected these in the fourth step of the data collection before including them in our analysis. Although there 69 errors in total in the data from their study, it is worth noting that we ran-ran the meta-analytical procedures from their study with the amended data and found that the major conclusions of their study were unaffected by these errors. In addition, we were thoroughly impressed by the amount of detail included in their supplementary files, and we were only able to check their data’s accuracy due to their diligence (cf. Ekroth et al. [9]). The different types of error made by the previous meta-analysis included (i) 27 comparisons that did not match the published raw data (available in the main text or online or in the supplementary material of each publication), (ii) 32 comparisons where effect sizes were calculated wrong and (iii) 10 comparisons which had not been transferred into the final metadata file correctly. There was one study which we could not check, because the original data was sent by personal communication from [31] to [10], nevertheless we included it in our analysis.
(4) *Extraction*: Fourth, for those studies or comparisons we had excluded (due to missing data or data errors) we extracted the data from the main text or supplementary files by going back to the original publications (we used PlotDigitizer (https://plotdigitizer.com/) to extract the information for any figures). In addition to the comparisons removed in the third step of data collection, this also included (i) 26 studies that, despite meeting our inclusion criteria, were removed because they were either missing the replicate-level raw data [9] or they were observational field studies based on multiple populations with a continuous measure of genetic diversity [10] and (ii) additional data for 3 comparisons [32–34] that were not made in the original analysis by Gibson & Nguyen [10].

We also collected information on 10 different moderators (see Table 1) by standardizing or recoding existing moderator variables used by Gibson & Nguyen [10], including host range, parasite diversity, metric of parasite success, host species, parasite type, source of host genetic diversity, scale of host diversity, mode of host reproduction, whether the parasite induces host mortality and whether the study was performed in a laboratory environment. Parasite diversity was not quantified as a continuous variable in the original studies, nor was it examined as part of the original experiments in most cases. Therefore, we binned parasite diversity into ‘high’ or ‘low’ groups depending on the following reasoning: ‘high parasite diversity’ if the isolate was collected from a natural population for a lab study, if the data was from an observational or experimental field study, or if more than one genotype had been identified (but this only applied to a small number of studies); ‘low parasite diversity’ if it was a laboratory strain, or only one genotype had been identified (but again, this only applied to a small number of studies). Where the information on these moderator variables was not already available from the supplementary material of Gibson & Nguyen [10] and was not available in the published article, we performed an online search to determine characteristics.

(5) *Recalculation*: Fifth, we calculated the mean, standard deviation and sample size for each comparison of high versus low genetic diversity groups of host populations for the data we extracted in the fourth step of data collection [10]. For certain studies, we calculated a pooled measure of the mean metric of parasite infection success for each group of high and low genetic diversity host populations, along with a pooled standard deviation and a pooled sample size. This included (i) studies with shared comparisons (e.g. multiple high diversity groups paired to the same low diversity group) [32,35] or (ii) those based on multiple populations with one or more continuous measures of genetic diversity [31, 34, 36–54]. In the latter case, the most appropriate measure of population-level genetic diversity was used (e.g. a measure of population-level genetic diversity based on Hardy-Weiberg equilibrium) and two separate groups of host and low diversity host populations were made with the same number of host populations in each group. In addition, host survival was converted into host mortality in some studies to reflect our definition of parasite infection success (see step two of data collection). Overall, this fifth step of data collection involved calculating parasite success data for 130 comparisons.

**Table 1.** Hypotheses for the influence of additional moderator variables on the nature of the effect of host population genetic diversity on mean and variability in parasite success.

| Moderator | Hypothesis |
| --- | --- |
| Metric of parasite success | Our study of 'parasite success' combined data of several types (eg prevalence, virulence, infection load). Using this moderator, we tested if the effects of host population genetic diversity differed between these different metrics. |
| Host type | The effect of host population genetic diversity may be influenced by the specificity of genetic interactions between host and parasite. These genetic interactions are thought to be more specific in invertebrates than in vertebrates [66], therefore we tested for inconsistency of the host population genetic diversity effect in these two groups. |
| Parasite type | Microparasites and macroparasites tend to have contrasting infection biology: varying in the duration of infections and the immune responses they trigger [67]. These differences might drive variation in the impact of host population genetic diversity. Therefore, we tested for inconsistency of the host population genetic diversity effect in these two groups. |
| Source of host genetic diversity | Studies typically investigate the impact of host genetic diversity by (i) controlling mating strategies to compare hosts with varying degrees of 'relatedness' (e.g. polyandry versus polygyny) or with different levels of inbreeding, (ii) using a suite of wildtype genotypes for controlled experiments with either low genetic diversity or high genetic diversity, or (iii) sampling organisms from the wild from populations that have been characterised as having different levels |
|  | of genetic diversity. We used this moderator to test if these different sources of genetic diversity affected the influence of host population genetic diversity. |
| Scale of host diversity | Host populations were predetermined as having either high or low diversity (discrete) or we separated them into such categories as part of our data collection (Fig 2, step 5) because the authors used multiple populations with a continuous measure of diversity. We used this moderator to test if this feature of how studies were designed had an effect on the influence of host population genetic diversity. |
| Mode of host reproduction | Host species reproduced sexually, asexually or using a combination of the two (i.e. facultatively sexual, such as *Daphnia*). We used this moderator to test if these different modes of host reproduction affected the influence of host population genetic diversity. |
| Host mortality? | The range of parasites studied can be further categorised by whether or not infection typically kills the host (which may be proxy for virulence). We used this moderator to test if differences in the virulent effects of parasitism affected the influence of host population genetic diversity. |
| Laboratory? | We used this moderator to test if differences in study setting (laboratory versus field) affected the influence of host population genetic diversity. |

After finishing all five steps of data collection, there was enough parasite success data to calculate both the SMD and lnCVR for 211 non-independent comparisons of high versus low genetic diversity groups of host populations.

### Calculation of effect sizes (SMD and lnCVR)

We calculated two effect size measures: the standardized mean difference (SMD) and the log coefficient of variation ratio (lnCVR). These quantified the effect of host population genetic diversity on either the mean or variability in parasite success across different studies [56]. This is because we were interested in comparing both the mean and variability in various measures of parasite infection success between groups of ‘high’ versus ‘low’ genetic diversity host populations.

We selected these specific ways of measuring between-group differences in either the mean or variability among metrics of parasite success because they took in to account certain factors: (i) SMD measures the mean difference between two groups (high versus low genetic diversity) in terms of standard deviations [56–57], so it can be used to compare metrics measured on very different scales (prevalence, load and virulence) [58]; it also corrects for small sample sizes, which is a common feature of ecological studies [59]. Similarly, lnCVR measures the ratio of variability between two groups adjusted for the size of the group means [60] and therefore can be used to compare multiple metrics. Also, it accounts for the possibility that the magnitude of the variability may scale with the mean, which is relevant for many types of count data that may be described by a Poisson distribution (e.g. parasite load).

Before calculating our effect sizes, we added a small value (0.001) to the mean and standard deviation in parasite success for each pair of high and low genetic diversity groups of host populations to ensure log values were calculated correctly. For consistency, we calculated SMD and its sampling variance from the formula derived from the supplementary material of [10], whereas we calculated variability effect sizes (lnCVR) and their sampling variances using the code from [60]. To account for comparisons based on a shared reference group of host populations with low genetic diversity (i.e. shared controls), we calculated the variance-covariance matrix for each effect size, using the make_VCV_matrix function from the metaAidR package v0.0.0.9000 [61].

All effect size calculations and subsequent calculations were performed in R v4.3.2 [62].

### Publication bias

Before analysing the data fully, we calculated the overall effect sizes for SMD and lnCVR and tested for any potential publication bias using funnel plots and Egger’s regression [63].

Meta-analytic models were fitted to the data using the rma.mv function from the metafor package v4.4.0 [64]. We included fixed effects for each type of effect size, the variance-covariance matrix of sampling errors, standard random effects for study and host genus, and correlated random effects for comparisons taken from the same experiment. Standard random effects for study and host genus were used to account for the possibility of non-independence between experiments originating from the same study and potential correlations between effects from closely related host species. Similarly, correlated random effects were used to account for potential non-independence of comparisons taken from the same experiment (multiple timepoints for a single comparison of a high and low genetic diversity group of host populations, or effect sizes from the same group of populations based on different measures of parasite success).

Funnel plots were used to identify whether published effect sizes were evenly distributed around model means by examining how outcomes varied as a function of their precision (standard error). Visual inspection of funnel plots for the effect of host population diversity on mean parasite success (Fig 3A) and its effect on the variability in parasite success (Fig 3B), showed no evidence for publication bias. More stringent evaluation showed that there was no correlation between the size of the effects themselves and their standard error (Egger’s test for both SMD and lnCVR: R = 0.06, 95% CI [-0.24, 0.37], p = 0.67 and R =-0.03, 95% CI [-0.38, 0.32], p = 0.86, respectively).

**Fig 3.**
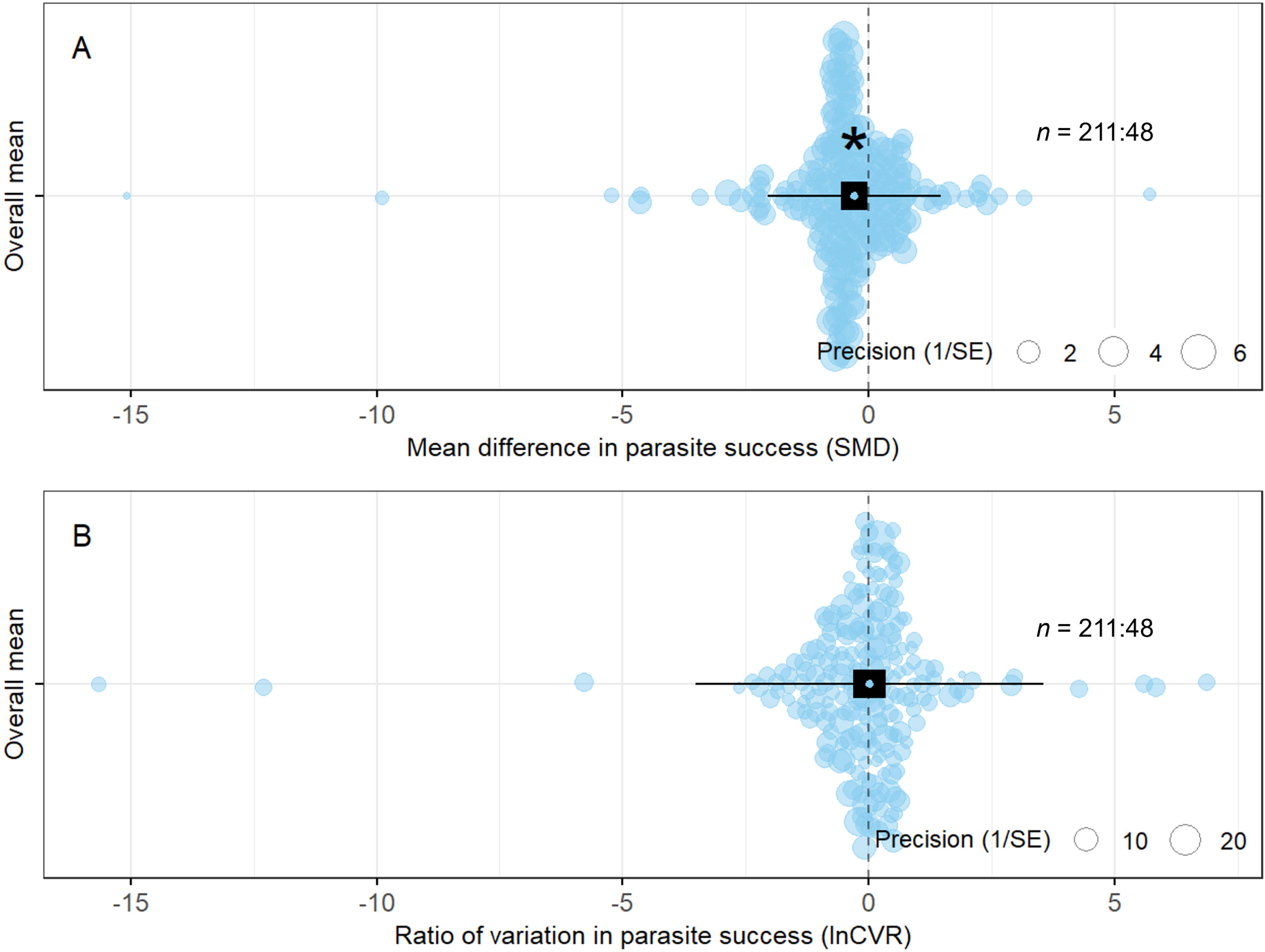
Testing for publication bias: the distribution of published effect sizes for our meta-analysis as a function of their precision (standard error). The x-axis in both plots shows effects of an increase in host population genetic diversity (high vs low) on A) the mean difference in parasite success (SMD) and B) the ratio of variation in parasite success (lnCVR). Model means and their 95% confidence intervals are shown by the dashed black lines.

### Meta-analysis of overall data

To test whether there was a significant difference in the mean parasite success (SMD) or variability in parasite success (lnCVR) between host populations with high versus low genetic diversity, we fitted mixed effects meta-analytic models. All of the models used in this paper were based on the same structure as those used for testing the presence of publication bias.

### Context dependence

*Moderators with an interaction-* To test if the overall effect of host population genetic diversity on the mean and variability in parasite success depended on an interaction between host range and parasite genetic diversity, we introduced an interaction term for these two moderators in our original meta-analytic models. Therefore, we could compare:

1) High versus low population genetic diversity of host specialist parasites.
2) High versus low population genetic diversity of host generalist parasites.

We compared the significance level of each individual predictor within the model, as well as the contrasts between them using the *glht* function from the *multcomp* package v1.4.25 [65].

*Moderators without an interaction –* To test our additional hypotheses (Table 1) for the eight remaining moderator variables, we modelled each moderator separately with its own individual mixed effects model. Before running the models, we removed redundant moderator categories with a limited sample size, such as transmission or infection rate (versus prevalence) and disease severity (versus load) for the metric of parasite success, and prokaryotic (versus vertebrate or invertebrate) for host species.

We compared the significance level of each individual predictor within the model, as well as the contrasts between them using ANOVA with a correction for multiple comparisons (Holm’s method).

### Sensitivity analysis of overall effects

To test the robustness of our results for the combined (overall) dataset, we performed a series of ‘leave-one-out’ sensitivity analyses. This involved the iterative exclusion of either one independent comparison (i.e. groups of high diversity host populations that shared a corresponding group of low diversity host populations were considered grouped together into a single comparison) or study at a time.

## Results

### Host population genetic diversity has an overall negative effect on mean parasite success

Our dataset contained 211 effect sizes from 48 different studies, including a range of (mainly animal) host and parasite species (Fig S2). These effect sizes represented individual estimates of the effect that change in host population genetic diversity has on parasite success; we assessed this effect on both mean parasite success (SMD) and the variability in parasite success (lnCVR).

Averaging over the whole data set, there was a significant effect of host population genetic diversity on mean parasite success (SMD =-0.29, 95% CI = [-0.57,-0.02], p = 0.04; Fig 4A): higher levels of host population genetic diversity were associated with lower mean parasite success. In contrast, across the whole data set, the variability of parasite success was not significantly affected by host population genetic diversity (lnCVR = 0.02, 95% CI = [-0.30, 0.35], p = 0.89; Fig 4B).

**Fig 4.**
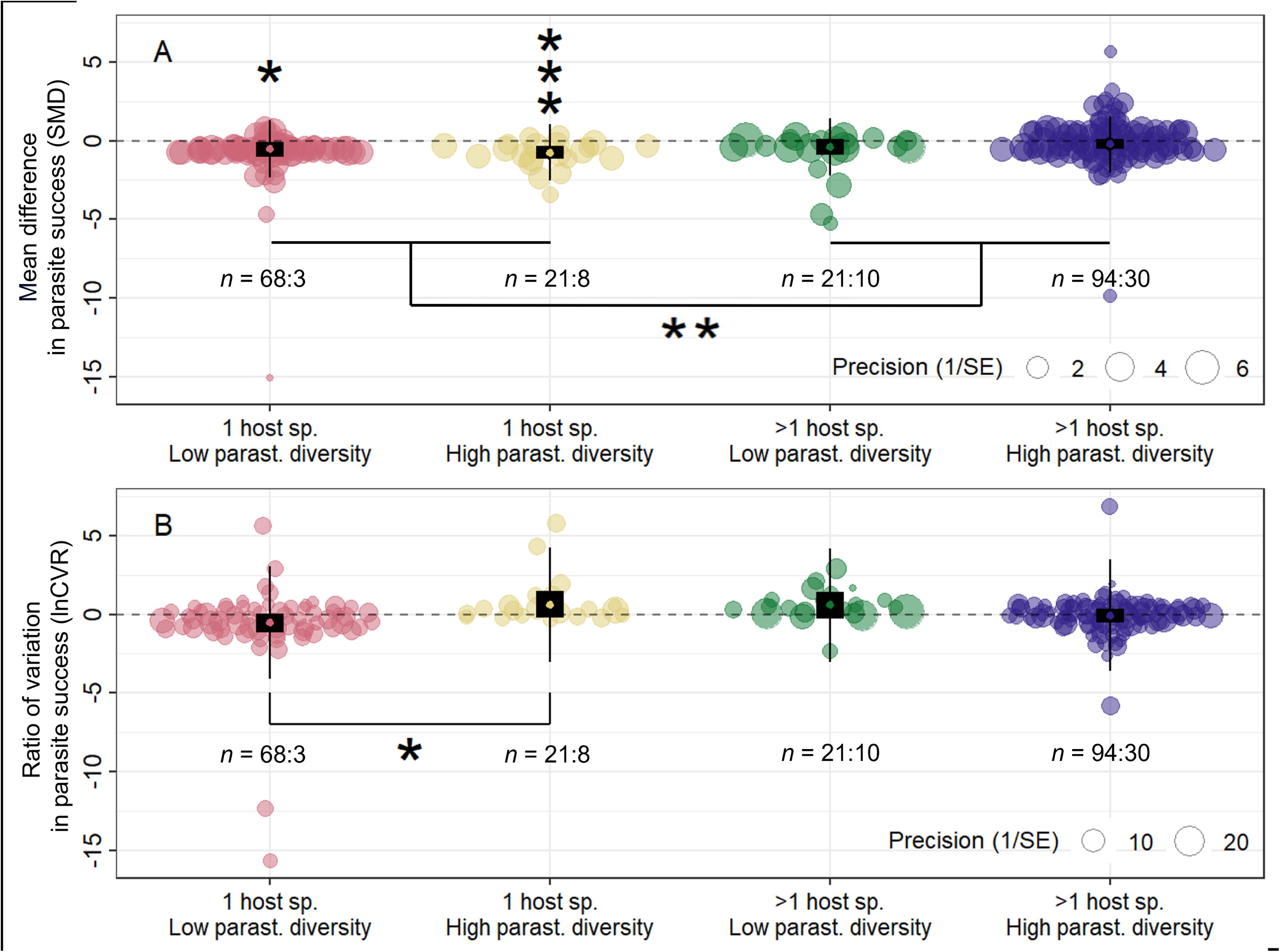
**The overall effect of host population genetic diversity on the mean and variability in parasite success**. The x-axis in each plot shows the effect that an increase in host population genetic diversity had on either A) the mean parasite success (SMD) or B) the variability in parasite success (lnCVR). The dashed line indicates an effect size of zero where host population genetic diversity has no influence. Model means are shown with 95% confidence intervals (black rectangles) and prediction intervals (thin black lines). Circles show individual effect sizes and are scaled according to the inverse of their standard error. n = sample size of the data (the number of effect sizes: the number of studies). The asterisk shows that the model mean is significantly different from zero (p < 0.05). Forest plot alternatives are shown in the online supplementary material (Fig S3).

In these analyses the residual variation (heterogeneity) in the data for both the difference in the mean and the variability of parasite success was high (I2 = 84.0% & 82.0% respectively). Most of this variation was explained by the effect of study (84.0% & 80.7%) and only a small amount was explained by host genus (0.0% & and 3.3%).

### Impacts of host population genetic diversity on parasite success differ between specialist and generalist parasites

Next, we investigated how the effect of host population genetic diversity on parasite infection success was influenced by two fundamental characteristics of the parasite: the host-specificity of the parasite and the likely genetic diversity of the parasite population studied.

The negative effect of host population genetic diversity on mean parasite infection success that we observed across the whole dataset (see above), was actually only significantly negative for specialist (single-host) parasites. This was true whether the observations for specialist parasites were for parasite populations with low or high genetic diversity (SMD = - 0.54, 95% CI = [-1.02,-0.06], p = 0.03 & SMD =-0.76, 95% CI = [-1.16,-0.37], p < 0.001 respectively; Fig 5A). The point estimates for the effect of host population genetic diversity on mean parasite infection success for generalist (multi-host) parasites were also negative; however, these estimates were not significant, and also not influenced by parasite isolate genetic diversity (SMD =-0.42, 95% CI = [-0.91, 0.07], p = 0.09 & SMD =-0.23, 95% CI = [-0.55, 0.08], p = 0.15 respectively; Fig 5A). This difference between specialist and generalist parasites in the effect of host population genetic diversity on mean parasite infection success was highly significant (glht = 0.47, SE = 0.16, p < 0.01; Fig 5A).

**Fig 5.**
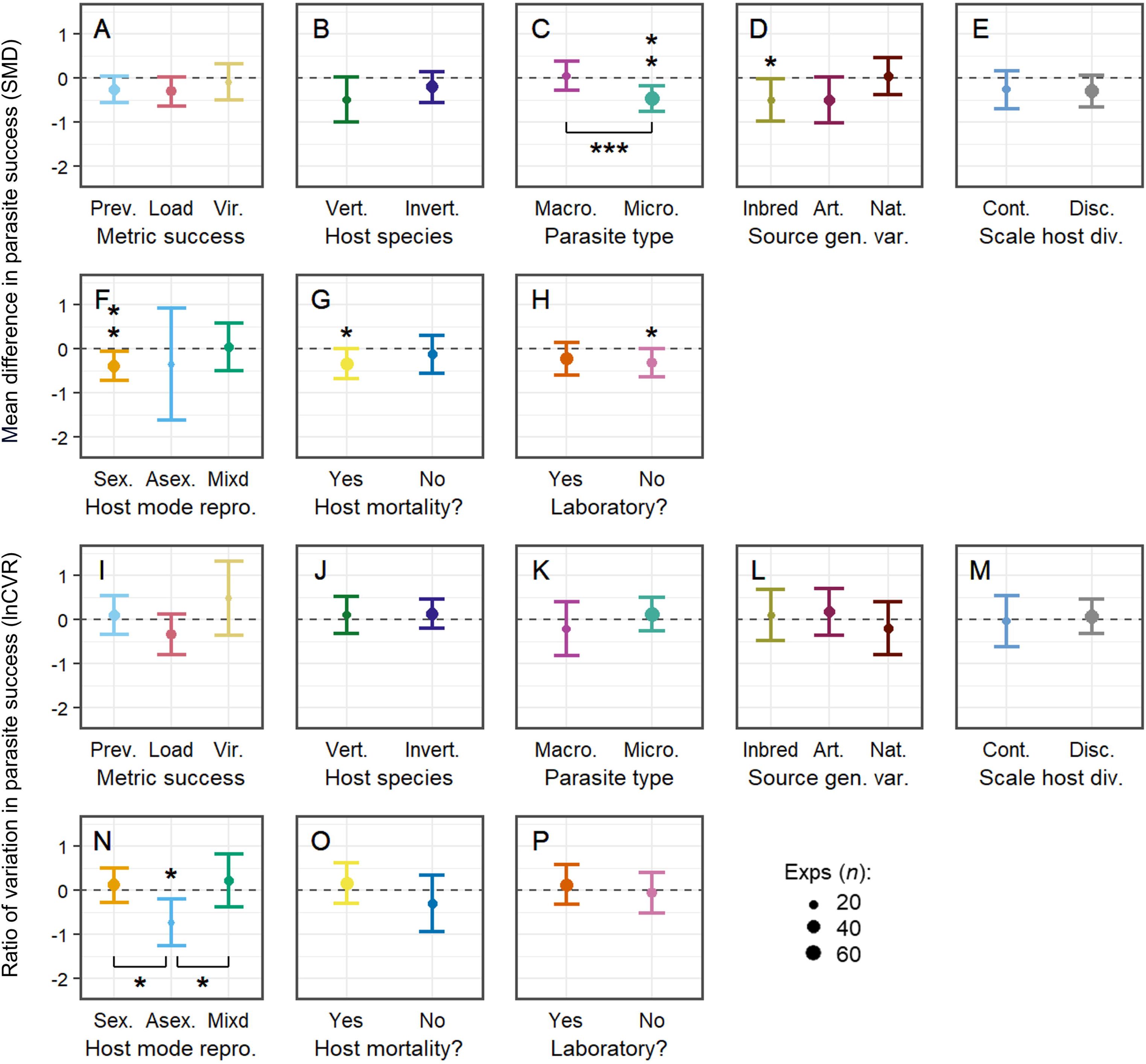
The effect of host population genetic diversity on both the mean and variability in metrics of parasite infection success depends on a combination of host range and parasite population genetic diversity. The x-axis in each plot shows the effect of an increase in host population genetic diversity on either A) mean parasite success (SMD) or B) variability in parasite success (lnCVR). The dashed line indicates an effect size of zero where there is no influence of host population genetic diversity on parasite success. Model means are shown with 95% confidence intervals (black rectangles) and prediction intervals (thin black lines). Individual effect sizes (circles) are scaled according to the inverse of their standard error. n = sample size of the data (the number of effect sizes: the number of studies). The significance level of individual model means, as well as any pairwise contrasts, is indicated by one (p < 0.05) or three (p < 0.001) asterisks.

Similar to the results for mean parasite success, the manner in which *variability* in parasite infection success was influenced by host population genetic diversity was also influenced by the characteristics of the parasite population. Although there was no significant relationship between host population genetic diversity and the variability in parasite success for the global dataset (see above), the observations for specialist parasites that infect only a single host species matched the predictions of our Epidemic Diversity conceptual model (Fig 5B).

When the genetic diversity of specialist parasites was low, an increase in host population genetic diversity tended to reduce variability in infection success (lnCVR =-0.54; 95% CI = [-1.14, 0.06], p = 0.08; Fig 5B). Whereas, for these specialist parasites, when parasite genetic diversity was high, an increase in host population genetic diversity resulted in increased variation in infection outcome (lnCVR = 0.61, 95% CI = [-0.23, 1.47], p = 0.16; Fig 5B). Although these associations were not significant in isolation, there was a significant difference between them (glht =-1.15, SE = 0.53 and p = 0.03). Also, mirroring our results for the mean of infection success, variability in parasite infection success was unaffected by host population genetic diversity in multi-host generalist parasites (Fig 5B).

### The effect of host population genetic diversity on parasite success is influenced by a number of other contextual factors

In addition to measuring the combined effect of parasite population genetic diversity and parasite host range on the relationship between host population genetic diversity and both the mean and variability in metrics of parasite infection success (see above), we also separately investigated how eight other contextual factors (Table 1) affected this relationship (Fig 6).

**Fig 6.** **The context-dependence of the effect of host population genetic diversity on the mean and variability in parasite success**. The y-axis in each plot shows the effect of an increase in host population genetic diversity on either the difference in mean parasite success (SMD) (panels A-H), or the difference in the variability in parasite success (lnCVR) (panels I-P). Model means are shown with 95% confidence intervals and are scaled according to the number of experiments. The dashed line indicates an effect size of zero.

Most of the moderator levels were not significantly different from zero and comparisons between different levels within the same moderator showed that they were often similar in terms of both the magnitude and direction of their effects. However, the parasite type (Fig 6C), source of host diversity (Fig 6D), host mode of reproduction (Fig 6F), whether the parasite kills the host or not (Fig 6G) and whether the study was performed in the laboratory (Fig 6H) all seemed to have a significant effect on the relationship between host population genetic diversity and mean parasite success. First, there was a significant reduction in mean microparasite success in host populations with higher genetic diversity (SMD =-0.47, 95% CI = [-0.77,-0.18], p < 0.01, Fig 6C) that was also associated with a significant difference between the effect of host population genetic diversity on microparasite versus macroparasite success (QM = 13.2, df = 1, p < 0.001, Fig 6C). Second, there was a significant reduction in mean parasite success for host populations with lower relatedness (outbred versus inbred) (SMD =-0.51, 95% CI = [-0.98,-0.03], p = 0.04, Fig 6D), similar to those composed of a higher number of chosen genotypes (close to the 95% significance threshold; SMD =-0.51, 95% CI = [-1.02, 0.01], p = 0.05, Fig 6D), but in contrast to host populations that were naturally more genetically diverse (SMD = 0.03, 95% CI = [-0.40, 0.45], p = 0.90, Fig 6D). Third, there was a significant reduction in mean parasite success in sexually reproducing host populations with higher genetic diversity (SMD =-0.40, 95% CI = [-073,-0.07], p = 0.02, Fig 6F) which was absent for both asexually and facultatively sexually reproducing host populations (SMD =-0.36, 95% CI = [-1.62, 0.91], p = 0.58 and SMD = 0.03, 95% CI = [-0.51, 0.57], p = 0.93 respectively, Fig 6F). Fourth, there was a significant reduction in mean parasite success for host populations with higher genetic diversity where the parasite usually killed the host (SMD =-0.34, 95% CI = [-0.68,-0.01], p < 0.05, Fig 6G) which was absent for parasites that did not usually kill the host (SMD =-0.13, 95% CI = - 0.56, 0.29], p = 0.54, Fig 6G). Fifth, there was a significant reduction in mean parasite success for host populations with higher genetic diversity studied outside of a laboratory environment (SMD =-0.33, 95% CI = [-0.65,-0.01], p = 0.04, Fig 6H) which was absent from studies conducted in a laboratory (SMD =-0.24, 95% CI = [-0.60, 0.13], p = 0.20, Fig 6H).

The significance level of individual model means, as well as any pairwise contrasts, is indicated by one (p < 0.05), two (p < 0.01) or three (p < 0.001) asterisks. The following abbreviations are used; Prev. (Prevalence), Vir. (Virulence), Vert. (Vertebrate), Invert. (Invertebrate), Macro. (Macroparasite), Micro. (Microparasite), Source gen. var. (Source of host genetic diversity), Art. (Artificial), Nat. (Natural), Scale host div. (Scale of host diversity), Cont. (Continuous), Disc. (Discrete), Host mode repro. (Mode of host reproduction), Sex. (Sexual), Asex. (Asexual), Mixd (Mixed).

There was only one moderator, the host mode of reproduction (Fig 6N), which had a significant effect on the relationship between host population genetic diversity and the variability in parasite success. Specifically, there was a significant reduction in the variability of parasite success for asexually reproducing host populations with higher genetic diversity (lnCVR =-0.74, 95% CI = [-1.26,-0.21], p < 0.01), Fig 6N) that was also associated with significant differences to the lack of any effect of host population genetic diversity on the variability in parasite success for both sexually and facultative-sexually reproducing hosts (QM = 6.40, df = 1, p = 0.01 & QM = 5.53, df = 1, p = 0.02 respectively).

### Some of our results are more robust than others

To test the robustness our of first analysis, which focussed on the overall effect of host population genetic diversity on parasite success, we performed a series of ‘leave-one-out’ sensitivity analyses. By averaging over multiple iterations, we showed that the overall effect of host population genetic diversity on both the mean and variability of parasite infection success was not dependent on the inclusion of any specific study or set of independent comparisons in our dataset. Specifically, the overall effect of host population genetic diversity on mean parasite infection success was significant across both sets of models (Table 2, Fig S4 and S5) and the overall effect of host population genetic diversity on variability in parasite infection success was still not significant (Table 2, Fig S4 and S5).

**Table 2.** Results of the leave-one-out sensitivity analyses.

| Method | ES | Estimate | SE | z-value | p-value | ci.lb | ci.ub |
| --- | --- | --- | --- | --- | --- | --- | --- |
| Leave1studyout | SMD | -0.29 | 0.14 | -2.05 | 0.04 | -0.57 | -0.01 |
| Leave1studyout | lnCVR | 0.02 | 0.17 | 0.13 | 0.87 | -0.31 | 0.35 |
| Leave1setout | SMD | -0.29 | 0.14 | -2.07 | 0.04 | -0.56 | -0.02 |
| Leave1setout | lnCVR | 0.02 | 0.17 | 0.13 | 0.9 | -0.3 | 0.35 |

Regarding the results of our main analysis, which focussed on how the mean and variability of parasite success was influenced by a three-way interaction between host population genetic diversity, parasite population genetic diversity and parasite host range, we were unable to perform any kind of formal sensitivity analysis due to the complexity of the underlying mixed-effects meta-analytical models. However, it is important to note that a large number of the effect sizes corresponding to the subset of data that included specialist parasites with low parasite population genetic diversity all came from a single study of a prokaryotic host (49 out of 68, [68]). Therefore, the results of some of these meta-analytical models may be quite sensitive to the inclusion of this one study.

Table 2 legend: Model averages were calculated from the iterative exclusion of either one study (Leave1studyout) or set of independent comparisons (Leave1setout). The following abbreviations are used; ES (effect size), SE (mean standard error across all models), ci.lb and ci.ub (mean lower and upper bounds of 95% confidence intervals across all models respectively).

## Discussion

By re-analysing the data from two previous studies [9,10] we show that the conventional theory that less genetically diverse host populations experience larger epidemics [6] is true only under certain circumstances. Despite finding a reduction in mean parasite infection success in host populations with higher genetic diversity that was consistent with previous studies [9,10], this result only applied to specialist parasites that infect only a single host species. In addition, we found support for our Epidemic Diversity conceptual model, which predicted that the *variability* in parasite success would depend on a combination of the population genetic diversity and host range of the parasite. For highly specific host-parasite interactions (as indicated by a narrow host range), we found that increased host population genetic diversity reduced the *variability* in parasite infection success when the corresponding parasite genetic diversity was low, but increased the *variability* in parasite infection success when parasite diversity was high. Therefore, the idea that less genetically diverse host populations generally experience larger epidemics may not be best described as ‘conventional wisdom’ [6].

Our results contrast those from previous meta-analytical studies of the effect of host population genetic diversity on metrics of parasite infection success, which also investigated the host range of the parasite as part of their analysis [9,10]. As already eluded to above, we found that the host range of the parasite was a significant moderator of the effect of host population genetic diversity on mean metrics of parasite infection success. Specifically, we found that a high level of host population genetic diversity tended to limit mean metrics of infection success for specialist, but not generalist, parasites relative to a low level of host population genetic diversity. One possible explanation for this is the increased statistical power of our study due to a larger number of effect sizes from combining the effect size data from these previous analyses [69]. This supports our original suggestion that parasite host range is closely related to the level of genetic specificity for infection (because specialist parasites are more likely to have evolved highly specific, matching-allele-type interactions between host resistance and parasite infectivity alleles than generalist parasites that are less tightly co-evolved to their range of hosts). It also highlights the susceptibility of host populations with only a small amount of genetic diversity to consistently high levels of infection success by specialist parasites.

Our finding that more diverse host populations tend to have smaller mean metrics of infection success for specialist parasites shows how there is slightly more complexity associated with conventional wisdom than previously thought [6]. It also has important implications for how the level of host population genetic diversity is managed in species of conservation concern [70]. For example, one approach to species management may be to prioritise the maintenance or restoration of genetic diversity in host populations threatened by specialist parasite species, or by finding a safe approach for broadening a specialist parasite’s host range. For example, the introduction of a novel host or parasite species, as some form of biological control [71], that can either act as a catalyst for host-mediated parasite evolution of greater generality [72] or cause a parasite host shift through direct competition for hosts (for a review, see [73]) may broaden the host range of a specialist parasite away from its target host to include a non-target, pest species. In addition, recent empirical work has started to test the theory that high host population genetic diversity (*sensu* ‘resource heterogeneity’) selects for the evolution, or maintenance, of a broader parasite host range (*sensu* ‘niche width’ [74]). Therefore, understanding how host population genetic diversity is linked to the evolution of parasite host range in a number of different host-parasite systems should be a priority for future research.

Again, in contrast to the results of previous meta-analytical studies [9,10], we also found that there was a significant difference between the effect of host population genetic diversity on the variability in metrics of infection success for specialist parasites with a high level of parasite population genetic diversity and a low level of parasite population genetic diversity. Specifically, we showed that a high level of both host and parasite population genetic diversity increased the variability in metrics of infection success for specialist parasites relative to host populations with a low level of population genetic diversity, whereas a high level of host population genetic diversity and a low level of parasite population genetic diversity decreased the variability in metrics of infections success for specialist parasites relative to host populations with a low level of population genetic diversity. Although these previous meta-analytical studies focused on the mean, rather than the variability in metrics of parasite infection success [9,10], nevertheless the authors of both studies had expected to find a significant effect of parasite population genetic diversity on the relationship between host population genetic diversity and mean metrics of parasite infection success and were surprised that there was no such significant result [9,10]. In addition to their reduced statistical power (as mentioned above), one possible reason for this could be that only one out of two of these studies investigated the interaction between different moderators [10]. On the other hand, the difference we observed in the variability in metrics of infection success for specialist parasites between host populations with high parasite population genetic diversity and low parasite population genetic diversity matched the initial predictions we made in our proposed Epidemic Diversity theoretical model (Fig 1). This confirms previous theories that the benefits of host population genetic diversity for resistance to disease depend on the corresponding parasite population genetic diversity [22,23,25].

This idea that the combination of both host and parasite population genetic diversity influence the variability in metrics of parasite infection success has important implications for host-parasite systems in general. As already mentioned previously, not only could such variability in metrics of parasite infection success be important for predicting the occurrence of potentially severe disease epidemics, which could benefit conservation by informing genetic diversity management strategies to prioritise at risk host populations [70], but it could also be central to our ability to protect against future emerging diseases [75] and for understanding the extent to which disease experiments are repeatable. For example, our results highlight that host populations with a low level of genetic diversity are particularly susceptible to consistently large disease epidemics caused by specialist parasites with a high level of diversity. Conversely, the inconsistent levels of parasite success predicted for combinations of low host x low parasite and high host x high parasite population genetic diversity suggest that the repeatability of both laboratory and field experiments may be quite low, since they are often characterised respectively by such combinations of host-parasite diversity. Similarly, patterns of future disease occurrence (and emergence) may be more difficult to predict in such systems compared to those with different combinations of diversity.

In addition to our moderator analysis using models with an interaction term, we also investigated the effects of eight other contextual factors to evaluate our list of hypotheses (Table 1). These are the same as the moderators used in previous meta-analyses [9,10], but in comparison to the total number of significant effects they observed in their analysis (two), our results show that there were six moderator levels that had significant effects. In particular, the effect of host population genetic diversity on mean metrics of infection success for microparasites was much more negative than for macroparasites. In agreement with our original hypothesis, this suggests that the difference in infection durability between micro-and macroparasites [67] affects the specificity of their interactions with the host [21]. Therefore, we suggest that macroparasites, due the longer-lasting nature of their infections [67], are less tightly coevolved with their hosts and thus have a lower genetic specificity for infection. We also found that there was a significant negative effect of host population genetic diversity on mean metrics of parasite infection success for comparisons of outbred versus inbred hosts. Although such an effect was absent for other host population comparisons, such as between naturally high and low genetic diversity populations of hosts, it was quite similar to the effect for host populations composed of select genotypes. Therefore, this could suggest that experimental manipulations of host population genetic diversity had a stronger effect on mean metrics of parasite infection success than studies using a purely natural source of hosts. However, it is worth noting that this result is somewhat inconsistent with the significantly negative effect of host population genetic diversity on mean metrics of parasite infection success observed for non-laboratory-based studies, for which the opposite effect was observed in one out of the two previous meta-analyses [9]. As such, an alternative explanation would be that the effect of host population genetic diversity on mean metrics of parasite infection success were exacerbated for outbred versus inbred hosts by the increased susceptibility of inbred hosts to disease [76].

Other notable observations from the individual models include a significant negative effect of host population genetic diversity on mean metrics of parasite infection success in sexually reproducing host populations and a significant negative effect of host population genetic diversity on the variability in metrics of parasite infection success for host populations reproducing asexually, which was strongly contrasted against the absence of either a sexually reproducing or facultatively sexually reproducing host. These results suggest that sexual reproduction might contribute to the strength of how population genetic diversity limits disease spread due to greater dissimilarity between genotypes from genetic recombination than achieved by asexual reproduction [77], but also that asexual reproduction can lead to greater disparity between the consistency of metrics of parasite infection success of host populations with high versus low genetic diversity than other forms of host reproduction. There was also a significant negative effect of host population genetic diversity on mean metrics of infection success for parasites that typically kill the host. Compared to less harmful parasites, this suggests that virulent parasites could select for higher levels of resistance and greater variation of resistance in the host population [9].

Despite the potentially exciting nature of our results, there are some additional considerations that should be taken into account. For example, there is a large number of effect sizes for specialist parasites with low population genetic diversity, but most of these actually come from a prokaryotic bacterial host study [68], rather than a vertebrate or invertebrate host, which is the case for most of our data. In addition, the host range of the parasite may not be a reliable estimate of the genetic specificity for infection. The host range of the parasite was used as a proxy for the genetic specificity for infection, as such a detailed level of information was not available. Therefore, we made the prediction that highly specific interactions between host and parasite genotypes [21] would be more likely for tightly coevolving pathogens (i.e. following a MAM of infection, [24]), as might be expected for specialist, but not generalist parasites. Similarly, the results of our moderator analysis rely on somewhat arbitrary ways of creating data sub-categories. In the case of parasite population genetic diversity, comparing mainly natural versus laboratory strains of parasites could be a poor indication of the effect of parasite population genetic diversity because the exact level of diversity was not actually quantified. In the case of the host range of the parasite, this measure is subjective and based somewhat on an incomplete literature [78].

One other final consideration is that the majority of our data concentrates on the effect of host population genetic diversity on both the mean and variability in metrics of parasite infection success for spatially replicated groups of host populations (but see [30]). Although we might expect the temporal pattern of the effect of host population genetic diversity on metrics of parasite infection success to be similar to that observed across space, we also predict some key differences. For example, recurrent bouts of parasite-mediated directional selection have the ability to reduce host and parasite population genetic diversity over time [79,80], which could be accompanied by a higher mean and lower variability in metrics of parasite infection success. However, the maintenance of host and parasite genetic diversity over time depends on the precise nature of selection and the underlying host-parasite infection genetics (i.e. a MAM versus a GFG model for genetic specificity, [81]). Although there are some studies which measure metrics of parasite infection success for host populations with different levels of genetic diversity at multiple timepoints (e.g. [29]), more studies would be required to provide a comprehensive test of the effect of host population genetic diversity on metrics of parasite infection success over time.

## Conclusions

In this study, we measured the difference in the mean and variability in metrics of parasite infection success between host populations with high versus low genetic diversity. After first challenging so-called ‘conventional wisdom’ (*sensu* [6]) we proposed an Epidemic Diversity model to better understand the context around how host population genetic diversity might affect not only the mean, but also the variability in metrics of parasite infection success. We found that host population genetic diversity affected mean metrics of infection success for specialist but not generalist parasites. We also found that the effect of host population diversity on the variability in metrics of parasite infection success depends on a combination of the host range of the parasite and the parasite population diversity, such that there is some evidence for an Epidemic Diversity theoretical model, at least for the collection of studies reviewed in this meta-analysis. Additionally, we found that there was a number of other context dependent effects of host population genetic diversity on both the mean and variability in metrics of parasite infection success, such as parasite type. Overall, these findings represent a change of perspective that could help to protect vulnerable host populations by prioritizing how genetic diversity within these populations is managed. Future study of the Epidemic Diversity hypothesis across a range of plant host-parasite systems would help generalize these findings.

## Data accessibility

Reviewer sharing

link: http://datadryad.org/share/LINK_NOT_FOR_PUBLICATION/E2NqLZ8KL2oYaPQLSYatmNNIeUub3aeC4ExfjJvu5Hw.

Data available from the Dryad Digital Repository: doi:10.5061/dryad.2bvq83bzq [82].

## Acknowledgements

We are grateful to Assistant Professor Alexander Strauss, Dr Camille Ameline, Dr Peter Thrall, Dr Jason Walsman and Assistant Professor Amanda Kyle Gibson for sharing their raw data and clarifying their methods with us. We also thank Dr Laura Braunholtz, Dr Katherine Raines, Dr Luc Bussière and Kyle Morrison for their general statistical and meta-analytical advice.

## Supporting information

**S1 Table. The difference between our amended study inclusion criteria and the original study inclusion criteria.** There is no S1 Table legend.

**S1 Fig. The difference between intra-and inter-group level variation in host population genetic diversity.** Each bucket represents a unique host population (e.g. jar of *Daphnia*). Each shape represents a unique host genotype. There are two sets of three host populations per replicate which correspond to the high and low diversity groups. The first two replicates show a combination of high intra-group variation in host population genetic diversity x low inter-group variation in host population genetic diversity. The second set of replicates show a combination of low intra-group variation in host population genetic diversity x high inter-group variation in host population genetic diversity.

**S2 Fig. The taxonomic distribution of unique host and parasite species (genera) within the data.** The number of unique combinations of host and parasite species are shown in each cell of the table, along with the number of studies they are sourced from after a backslash. The total number of studies (59) is higher than the total number of studies in our dataset (48), because there were some studies with multiple comparisons of unique host and parasite combinations. The colour system corresponds to the number of unique combinations of host and parasite genera, where higher numbers have darker colouration.

**S3 Fig. Study effects of host population genetic diversity on the mean and variability in parasite success.** The x-axis in each plot shows the effect of increasing host population genetic diversity on either A) the difference in mean parasite success (SMD) or B) the difference in the variability in parasite success (lnCVR). Aggregated effects for each study are shown with 95% confidence intervals. Where the same host genus was studied more than once (‘Duplicated Genus’), the colour of the points is white, rather than black, and the specific host genus studied is indicated by its shape (there were only five duplicated host genera). Each point is scaled by the amount of weighting they received in an aggregated mixed effects model, whereas the actual analysis was conducted based on the full set of 211 individual data points. The dashed lines indicate an effect size of zero and the overall model means are shown by the solid grey line with 95% confidence intervals bands in light grey.

**S4 Fig. The results of the leave-one-study-out method of sensitivity analysis.** The x-axis in each plot shows the effect of increasing host population genetic diversity on either A) the difference in mean parasite success (SMD) or B) the difference in the variability in parasite success (lnCVR). The names of the authors and the publication date for the study omitted in each model iteration is shown on the left, with the overall effect size and its confidence interval shown on in the middle. The meta-regression estimate of the original model using the full set of studies is shown by the vertical line and the specific value for each individual study is shown on the right (with 95% confidence intervals). The size of each point is scaled according to its precision.

**S5 Fig. The results of the leave-one-independent-comparison-out method of sensitivity analysis visualized using a modified version of an orchard plot.** The x-axis in each plot shows the effect of increasing host population genetic diversity on either A) the difference in mean parasite success (SMD) or B) the difference in the variability in parasite success (lnCVR). Unlike traditional orchard plots, which show the distribution of individual effect sizes, the mean effect size for each model iteration is shown by the coloured circles.

The size of each point is scaled by its precision (inverse of the standard error). The meta-regression estimate of the original model using the full set of studies is shown by the dotted line.

## References

1. Dicker RC, Coronado F, Koo D, Parrish RG. Principles of epidemiology in public health practice; an introduction to applied epidemiology and biostatistics. 3rd ed. Atlanta: U.S. Department of Health and Human Services; 2006.

2. Altizer S, Harvell D, Friedle E. Rapid evolutionary dynamics and disease threats to biodiversity. Trends Ecol Evol. 2003;18: 589–596. doi:10.1016/J.TREE.2003.08.013

3. Zhu Y, Chen H, Fan J, Wang Y, Li Y, Chen J, et al. Genetic diversity and disease control in rice. Nature. 2000;406: 718–722. doi:10.1038/35021046

4. De Castro F, Bolker B. Mechanisms of disease-induced extinction. Ecol Lett. 2005;8: 117–126. doi:10.1111/j.1461-0248.2004.00693.x

5. Jones KE, Patel NG, Levy MA, Storeygard A, Balk D, Gittleman JL, et al. Global trends in emerging infectious diseases. Nature. 2008;451: 990–993. doi:10.1038/nature06536

6. King KC, Lively CM. Does genetic diversity limit disease spread in natural host populations? Heredity. 2012;109: 199–203. doi:10.1038/hdy.2012.33

7. Elton CS. The Ecology of Invasions by Animals and Plants. Boston, MA: Springer US; 1958. doi:10.1007/978-1-4899-7214-9

8. Garrett KA, Mundt CC. Epidemiology in mixed host populations. Phytopathology. 1999;89: 984–990. doi:10.1094/PHYTO.1999.89.11.984

9. Ekroth AKE, Rafaluk-Mohr C, King KC. Host genetic diversity limits parasite success beyond agricultural systems: a meta-analysis. Proceedings of the Royal Society B: Biological Sciences. 2019;286: 20191811. doi:10.1098/rspb.2019.1811

10. Gibson AK, Nguyen AE. Does genetic diversity protect host populations from parasites? A meta-analysis across natural and agricultural systems. Evol Lett. 2020;5: 16–32. doi:10.1002/evl3.206

11. Altizer S, Hochachka WM, Dhondt AA. Seasonal dynamics of mycoplasmal conjunctivitis in eastern North American house finches. Journal of Animal Ecology. 2004. doi:10.1111/j.0021-8790.2004.00807.x

12. Aznar MN, Linares FJ, Cosentino B, Sago A, La Sala L, León E, et al. Prevalence and spatial distribution of bovine brucellosis in San Luis and La Pampa, Argentina. BMC Vet Res. 2015;11. doi:10.1186/s12917-015-0535-1

13. Cáceres CE, Hall SR, Duffy MA, Tessier AJ, Helmle C, MacIntyre S. Physical structure of lakes constrains epidemics in Daphnia populations. Ecology. 2006;87: 1438–1444. doi:10.1890/0012-9658(2006)87[1438:PSOLCE]2.0.CO;2

14. Carlsson-Granér U, Thrall PH. The spatial distribution of plant populations, disease dynamics and evolution of resistance. Oikos. 2002;97. doi:10.1034/j.1600-0706.2002.970110.x

15. Ericson L, Burdon JJ, Müller WJ. Spatial and temporal dynamics of epidemics of the rust fungus Uromyces valerianae on populations of its host Valeriana salina. Journal of Ecology. 1999;87. doi:10.1046/j.1365-2745.1999.00384.x

16. Montano S, Giorgi A, Monti M, Seveso D, Galli P. Spatial variability in distribution and prevalence of skeletal eroding band and brown band disease in Faafu Atoll, Maldives. Biodivers Conserv. 2016;25. doi:10.1007/s10531-016-1145-3

17. Thrall PH, Laine A-L, Ravensdale M, Nemri A, Dodds PN, Barrett LG, et al. Rapid genetic change underpins antagonistic coevolution in a natural host-pathogen metapopulation. Ecol Lett. 2012;15: 425–435. doi:10.1111/j.1461-0248.2012.01749.x

18. Vergara D, Lively CM, King KC, Jokela J. The geographic mosaic of sex and infection in lake populations of a New Zealand snail at multiple spatial scales. American Naturalist. 2013;182. doi:10.1086/671996

19. Tarpy DR. Genetic diversity within honeybee colonies prevents severe infections and promotes colony growth. Proceedings of the Royal Society B: Biological Sciences. 2003;270. doi:10.1098/rspb.2002.2199

20. Paplauskas S, Brand J, Auld SKJR. Ecology directs host–parasite coevolutionary trajectories across Daphnia–microparasite populations. Nat Ecol Evol. 2021;5: 480–486. doi:10.1038/s41559-021-01390-7

21. Schmid-Hempel P, Ebert D. On the evolutionary ecology of specific immune defence. Trends Ecol Evol. 2003;18: 27–32. doi:10.1016/S0169-5347(02)00013-7

22. Boomsma JJ. Paternity in eusocial Hymenoptera. Philos Trans R Soc Lond B Biol Sci. 1996;351: 947–975. doi:10.1098/rstb.1996.0087

23. Van Baalen M, Beekman M. The costs and benefits of genetic heterogeneity in resistance against parasites in social insects. American Naturalist. 2006. doi:10.1086/501169

24. Agrawal A, Lively CM. Infection genetics: Gene-for-gene versus matching-alleles models and all points in between. Evol Ecol Res. 2002;4: 79–90.

25. Bensch HM, O’Connor EA, Cornwallis CK. Living with relatives offsets the harm caused by pathogens in natural populations. Elife. 2021;10. doi:10.7554/eLife.66649

26. Luijckx P, Fienberg H, Duneau D, Ebert D. A matching-allele model explains host resistance to parasites. Current Biology. 2013;23: 1085–1088. doi:10.1016/j.cub.2013.04.064

27. Reiss ER, Drinkwater LE. Cultivar mixtures: A meta-analysis of the effect of intraspecific diversity on crop yield: A. Ecological Applications. 2018. doi:10.1002/eap.1629

28. Baer B, Schmid-Hempel P. Experimental variation in polyandry affects parasite loads and fitness in a bumble-bee. Nature. 1999;397: 151–154. doi:10.1038/16451

29. Altermatt F, Ebert D. Genetic diversity of Daphnia magna populations enhances resistance to parasites. Ecol Lett. 2008;10: 44–53. doi:10.1111/j.1461-0248.2006.00995.x

30. Hale KA, Briskie J V. Decreased immunocompetence in a severely bottlenecked population of an endemic New Zealand bird. Anim Conserv. 2007;10. doi:10.1111/j.1469-1795.2006.00059.x

31. King KC, Jokela J, Lively CM. Parasites, sex, and clonal diversity in natural snail populations. Evolution. 2011;65. doi:10.1111/j.1558-5646.2010.01215.x

32. Agha R, Gross A, Rohrlack T, Wolinska J. Adaptation of a Chytrid Parasite to Its Cyanobacterial Host Is Hampered by Host Intraspecific Diversity. Front Microbiol. 2018;9: 1–10. doi:10.3389/fmicb.2018.00921

33. Baer B, Schmid-Hempel P. Unexpected consequences of polyandry for parasitism and fitness in the bumblebee, Bombus terrestris. Evolution. 2001. doi:10.1111/j.0014-3820.2001.tb00683.x

34. Giese AR, Hedrick PW. Genetic variation and resistance to a bacterial infection in the endangered Gila topminnow. Anim Conserv. 2003;6. doi:10.1017/S1367943003003445

35. Schmidt AM, Linksvayer TA, Boomsma JJ, Pedersen JS. No benefit in diversity? The effect of genetic variation on survival and disease resistance in a polygynous social insect. Ecol Entomol. 2011;36. doi:10.1111/j.1365-2311.2011.01325.x

36. Dagan Y, Liljeroos K, Jokela J, Ben-Ami F. Clonal diversity driven by parasitism in a freshwater snail. J Evol Biol. 2013;26. doi:10.1111/jeb.12245

37. Dionne M, Miller KM, Dodson JJ, Bernatchez L. MHC standing genetic variation and pathogen resistance in wild Atlantic salmon. Philosophical Transactions of the Royal Society B: Biological Sciences. 2009;364. doi:10.1098/rstb.2009.0011

38. Ellison A, Cable J, Consuegra S. Best of both worlds? Association between outcrossing and parasite loads in a selfing fish. Evolution. 2011;65: 3021–3026. doi:10.1111/j.1558-5646.2011.01354.x

39. Field SG, Lange M, Schulenburg H, Velavan TP, Michiels NK. Genetic diversity and parasite defense in a fragmented urban metapopulation of earthworms. Anim Conserv. 2007;10. doi:10.1111/j.1469-1795.2006.00084.x

40. Kyle CJ, Rico Y, Castillo S, Srithayakumar V, Cullingham CI, White BN, et al. Spatial patterns of neutral and functional genetic variations reveal patterns of local adaptation in raccoon (Procyon lotor) populations exposed to raccoon rabies. Mol Ecol. 2014;23. doi:10.1111/mec.12726

41. Loiseau C, Zoorob R, Robert A, Chastel O, Julliard R, Sorci G. Plasmodium relictum infection and MHC diversity in the house sparrow (Passer domesticus). Proceedings of the Royal Society B: Biological Sciences. 2011;278. doi:10.1098/rspb.2010.1968

42. Meagher S. Genetic diversity and capillaria hepatica (nematoda) prevelance in Michigan deer mouse populations. Evolution. 1999;53: 1318–1324. doi:10.2307/2640837

43. Neumann P, Moritz RFA. Testing genetic variance hypotheses for the evolution of polyandry in the honeybee (Apis mellifera L.). Insectes Soc. 2000;47. doi:10.1007/PL00001714

44. Parsche S, Lattorff MHG. The relative contributions of host density and genetic diversity on prevalence of a multi-host parasite in bumblebees. Biological Journal of the Linnean Society. 2018;125. doi:10.1093/biolinnean/bly151

45. Pierce AA, De Roode JC, Altizer S, Bartel RA. Extreme heterogeneity in parasitism despite low population genetic structure among monarch butterflies inhabiting the Hawaiian Islands. PLoS One. 2014;9. doi:10.1371/journal.pone.0100061

46. Puurtinen M, Hytönen M, Knott KE, Taskinen J, Nissinen K, Kaitala V. The effects of mating system and genetic variability on susceptibility to trematode parasites in a freshwater snail, Lymnaea stagnalis. Evolution. 2004;58. doi:10.1111/j.0014-3820.2004.tb01626.x

47. Queirós J, Vicente J, Alves PC, de la Fuente J, Gortazar C. Tuberculosis, genetic diversity and fitness in the red deer, Cervus elaphus. Infection, Genetics and Evolution. 2016;43. doi:10.1016/j.meegid.2016.05.031

48. Rahn AK, Krassmann J, Tsobanidis K, MacColl ADC, Bakker TCM. Strong neutral genetic differentiation in a host, but not in its parasite. Infection, Genetics and Evolution. 2016;44. doi:10.1016/j.meegid.2016.07.011

49. Savage AE, Becker CG, Zamudio KR. Linking genetic and environmental factors in amphibian disease risk. Evol Appl. 2015;8. doi:10.1111/eva.12264

50. Trouvé S, Degen L, Renaud F, Goudet J. Evolutionary implications of a high selfing rate in the freshwater snail Lymnaea Truncatula. Evolution. 2003;57. doi:10.1554/02-452

51. Velavan TP, Weller S, Schulenburg H, Michiels NK. High genetic diversity and heterogeneous parasite load in the earthworm Lumbricus terrestris on a German meadow. Soil Biol Biochem. 2009;41. doi:10.1016/j.soilbio.2009.03.026

52. Whitehorn PR, Tinsley MC, Brown MJF, Darvill B, Goulson D. Genetic diversity, parasite prevalence and immunity in wild bumblebees. Proceedings of the Royal Society B: Biological Sciences. 2011;278: 1195–1202. doi:10.1098/rspb.2010.1550

53. Whitehorn PR, Tinsley MC, Brown MJF, Darvill B, Goulson D. Genetic diversity and parasite prevalence in two species of bumblebee. J Insect Conserv. 2014;18. doi:10.1007/s10841-014-9673-1

54. Whiteman NK, Matson KD, Bollmer JL, Parker PG. Disease ecology in the Galápagos Hawk (Buteo galapagoensis): host genetic diversity, parasite load and natural antibodies. Proceedings of the Royal Society B: Biological Sciences. 2006;273: 797–804. doi:10.1098/rspb.2005.3396

55. Page MJ, McKenzie JE, Bossuyt PM, Boutron I, Hoffmann TC, Mulrow CD, et al. The PRISMA 2020 statement: an updated guideline for reporting systematic reviews. BMJ. 2021; n71. doi:10.1136/bmj.n71

56. Borenstein M, Hedges L V., Higgins JPT, Rothstein HR. Introduction to meta-analysis. Introduction to Meta-Analysis. 2009. doi:10.1002/9780470743386

57. Field AP, Gillett R. How to do a meta-analysis. British Journal of Mathematical and Statistical Psychology. 2010;63. doi:10.1348/000711010X502733

58. Higgins JPT, Green S. Cochrane handbook for systematic reviews of interventions. The Cochrane Collaboration; 2024. Available: www.handbook.cochrane.org

59. Jennions MD. A survey of the statistical power of research in behavioral ecology and animal behavior. Behavioral Ecology. 2003;14: 438–445. doi:10.1093/beheco/14.3.438

60. Nakagawa S, Poulin R, Mengersen K, Reinhold K, Engqvist L, Lagisz M, et al. Meta-analysis of variation: ecological and evolutionary applications and beyond. O’ Hara RB, editor. Methods Ecol Evol. 2015;6: 143–152. doi:10.1111/2041-210X.12309

61. Lagisz M, Senior A, Nakagawa S, Noble D. _metaAidR: Functions and Data for Aiding Meta-analyses in Ecology and Evolution_. 2024.

62. R Core Team. _R: A Language and Environment for Statistical Computing_. Vienna, Austria: R Foundation for Statistical Computing; 2023. Available: https://www.r-project.org/

63. Sutton AJ. Publication bias. In: Cooper H, Hedges L V, Valentine JC, editors. The handbook of research synthesis and meta-analysis. Russell Sage Foundation; 2009. pp. 435–452.

64. Viechtbauer W. Conducting Meta-analysis in R with the metafor package. J Stat Softw. 2010;36.

65. Hothorn T, Bretz F, Westfall P. Simultaneous inference in general parametric models. Biometrical Journal. 2008;50: 346–363. doi:10.1002/bimj.200810425

66. Dybdahl MF, Jenkins CE, Nuismer SL. Identifying the molecular basis of host-parasite coevolution: Merging models and mechanisms. Am Nat. 2014;184: 1–13. doi:10.1086/676591

67. Sorci G. Immunology of parasitism. Reference Module in Biomedical Sciences. Elsevier; 2014. doi:10.1016/B978-0-12-801238-3.00253-1

68. Van Houte S, Ekroth AKE, Broniewski JM, Chabas H, Ashby B, Bondy-Denomy J, et al. The diversity-generating benefits of a prokaryotic adaptive immune system. Nature. 2016;532. doi:10.1038/nature17436

69. Gibson AK. Genetic diversity and disease: The past, present, and future of an old idea. Evolution. 2022;76: 20–36. doi:10.1111/evo.14395

70. Meuwissen THE, Sonesson AK, Gebregiwergis G, Woolliams JA. Management of genetic diversity in the era of genomics. Front Genet. 2020;11. doi:10.3389/fgene.2020.00880

71. Stenberg JA, Sundh I, Becher PG, Björkman C, Dubey M, Egan PA, et al. When is it biological control? A framework of definitions, mechanisms, and classifications. J Pest Sci. 2021;94: 665–676. doi:10.1007/s10340-021-01354-7

72. Bull JJ, Wichman HA, Krone SM. Modeling the directed evolution of broad host range Phages. Antibiotics. 2022;11: 1709. doi:10.3390/antibiotics11121709

73. Bashey F. Within-host competitive interactions as a mechanism for the maintenance of parasite diversity. Philosophical Transactions of the Royal Society B: Biological Sciences. 2015;370: 20140301. doi:10.1098/rstb.2014.0301

74. Gibson AK, Baffoe-Bonnie H, Penley MJ, Lin J, Owens R, Khalid A, et al. The evolution of parasite host range in heterogeneous host populations. J Evol Biol. 2020;33: 773–782. doi:10.1111/jeb.13608

75. Altizer S, Dobson A, Hosseini P, Hudson P, Pascual M, Rohani P. Seasonality and the dynamics of infectious diseases. Ecol Lett. 2006;9: 467–484. doi:10.1111/j.1461-0248.2005.00879.x

76. Coltman DW, Pilkington JG, Smith JA, Pemberton JM. Parasite-medidated selection against inbred soay sheep in a free-living, island population. Evolution. 1999. Available: https://about.jstor.org/terms

77. Hamilton WD, Axelrod R, Tanese R. Sexual reproduction as an adaptation to resist parasites (A review). Proc Natl Acad Sci U S A. 1990;87: 3566–3573. doi:10.1073/pnas.87.9.3566

78. Hyman P, Abedon ST. Bacteriophage host range and bacterial resistance. Adv Appl Microbiol. 2010;70: 217–48. doi:10.1016/S0065-2164(10)70007-1

79. Buckling A, Rainey PB. Antagonistic coevolution between a bacterium and a bacteriophage. Proceedings of the Royal Society B: Biological Sciences. 2002;269: 931–936. doi:10.1098/rspb.2001.1945

80. Obbard DJ, Jiggins FM, Bradshaw NJ, Little TJ. Recent and recurrent selective sweeps of the antiviral RNAi gene Argonaute-2 in three species of Drosophila. Mol Biol Evol. 2011;28: 1043–56. doi:10.1093/molbev/msq280

81. Boots M, White A, Best A, Bowers R. How specificity and epidemiology drive the coevolution of static trait diversity in hosts and parasites. Evolution. 2014;68: 1594– 1606. doi:10.1111/evo.12393

82. Paplauskas S, Duthie B, Tinsley MC. An ‘Epidemic Diversity’ conceptual model explains how host genetic diversity affects variation in parasite success [Dataset]. Dryad. 2025. 10.5061/dryad.2bvq83bzq

