## Supplementary figures and images for "An ‘Epidemic Diversity’ conceptual model explains how host genetic diversity affects variation in parasite success"

### Supplemental Figure 1

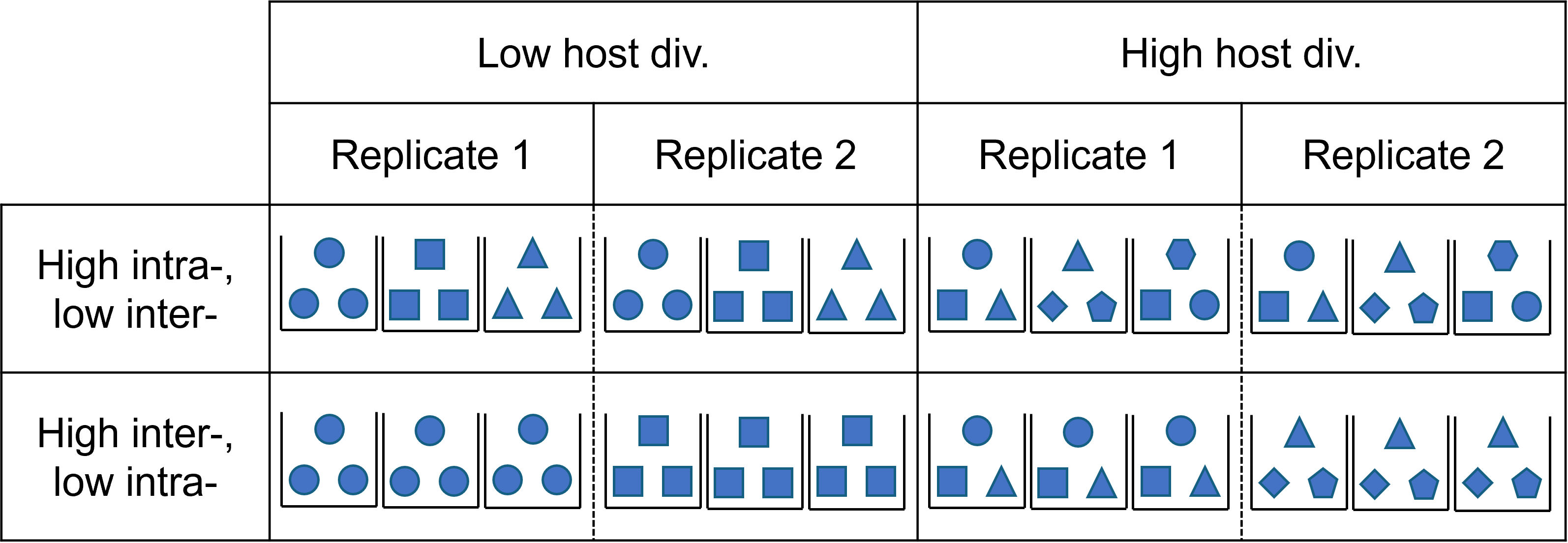

### Supplemental Figure 2

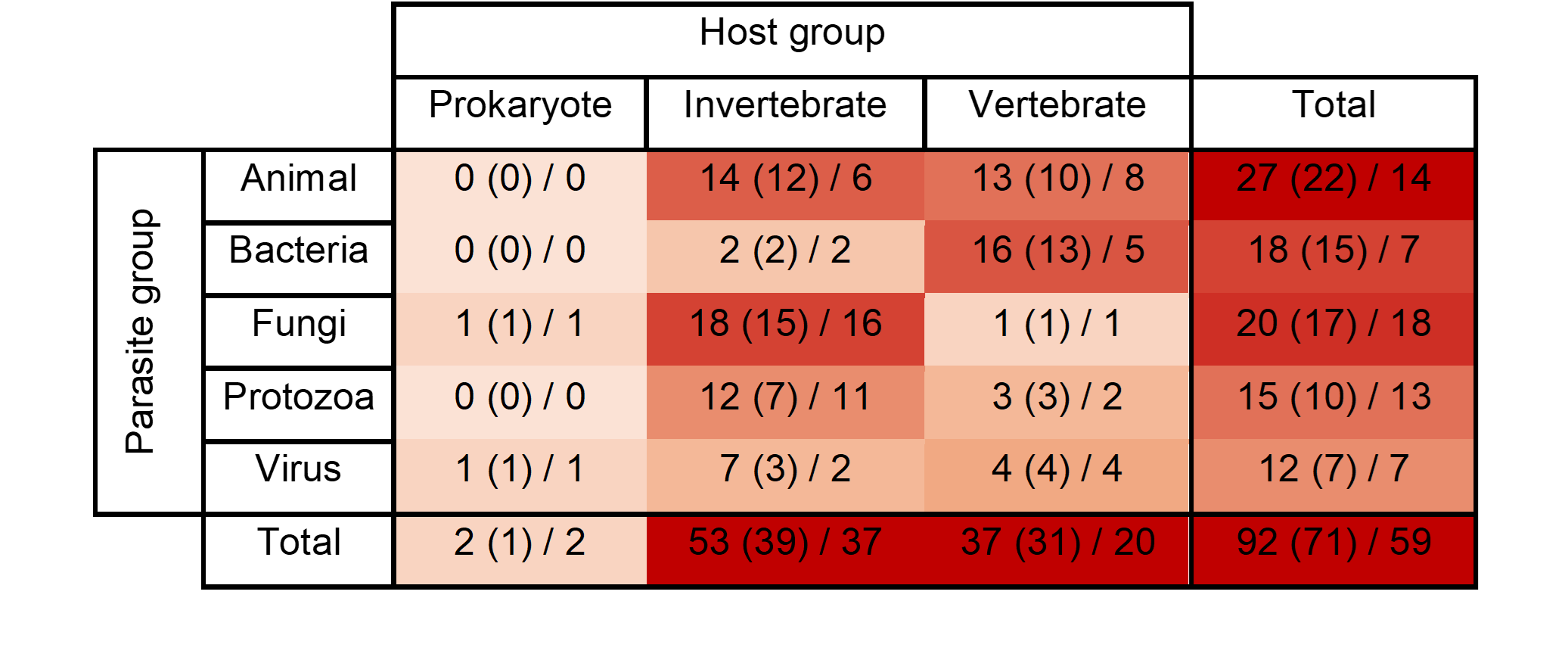

### Supplemental Figure 3

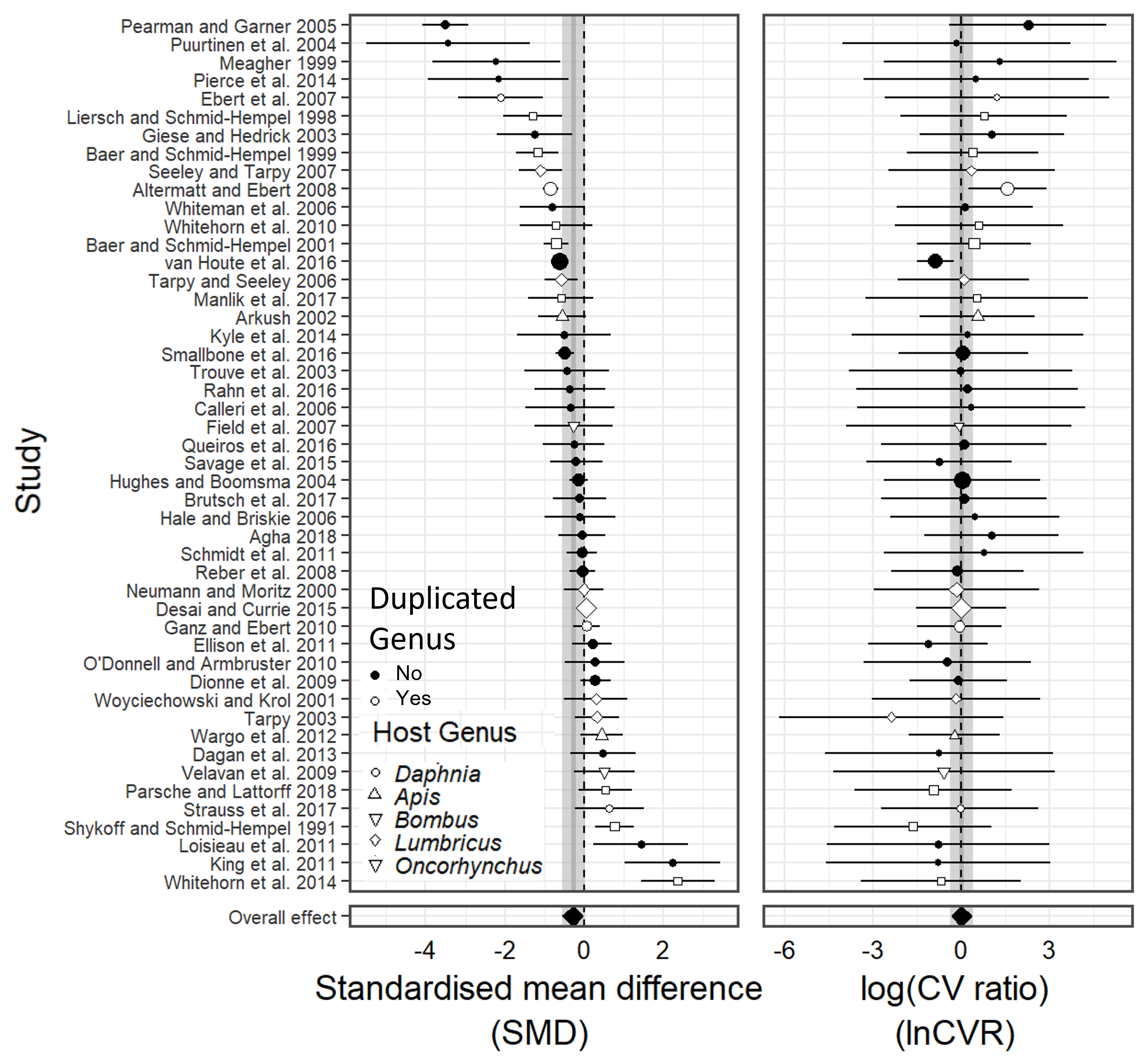

### Supplemental Figure 4

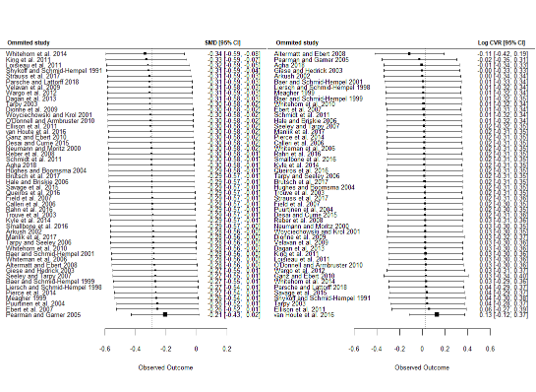

### Supplemental Figure 5

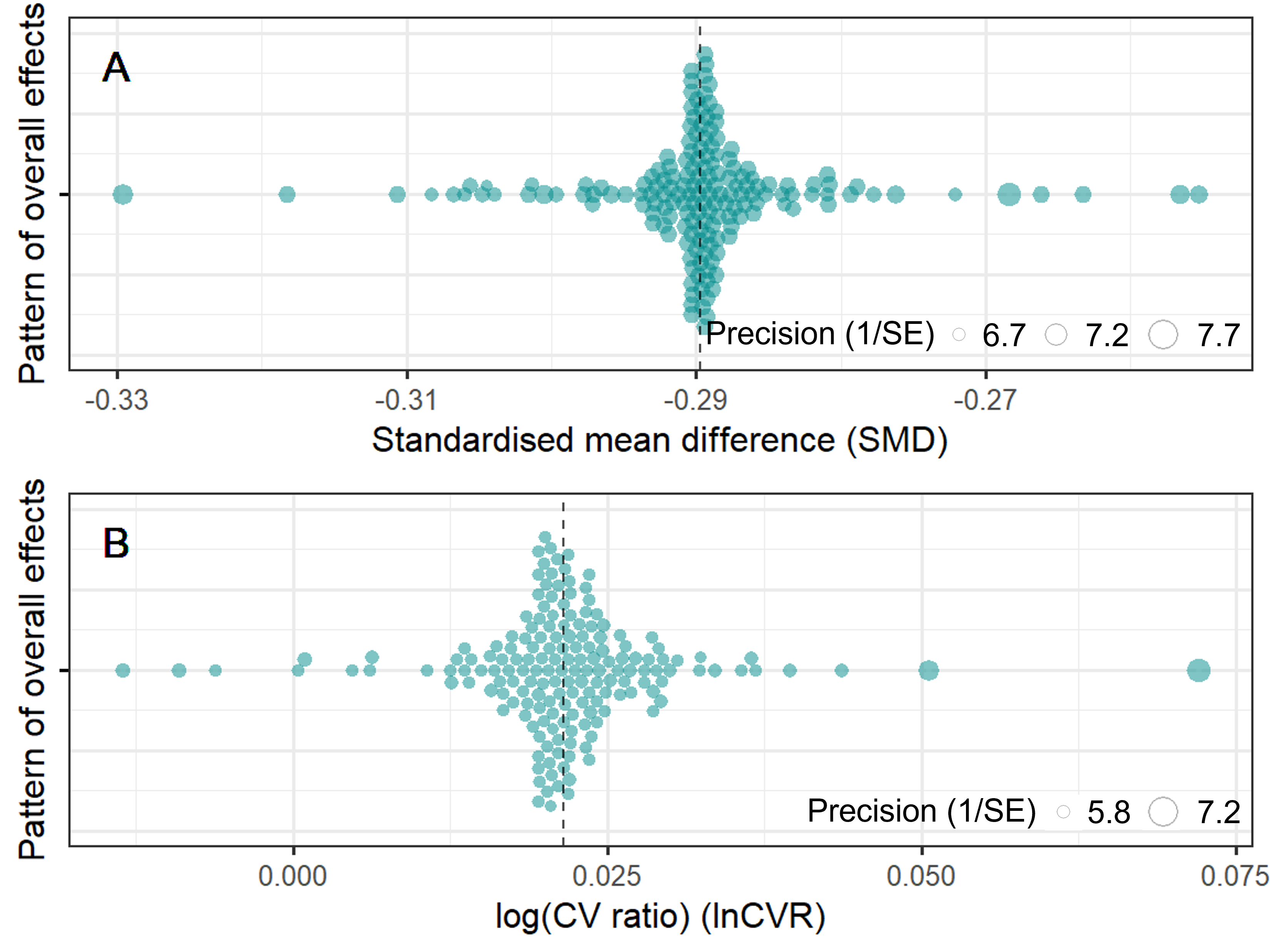
