## Supplemental Table 1 for "An ‘Epidemic Diversity’ conceptual model explains how host genetic diversity affects variation in parasite success"

**S1 Table. The difference between our amended study inclusion criteria and the original study inclusion criteria.**

| New study inclusion criteria | Original study inclusion criteria | Why changed |
| --- | --- | --- |
| 1) ‘Parasite success’, which we define as the ability of a parasite to spread among hosts (transmission rate, infection rate, prevalence), replicate on / within hosts (macro / microparasite load, disease severity), or kill hosts (virulence i.e. host survival / mortality rate) was measured among replicate populations across time or space. | - Ekroth, Rafaluk-Mohr and King, 2019: Defined parasite success as any measure of a parasite’s ability to proliferate within a host population.  - Gibson and Nguyen, 2020: Focused on population-level parasitism, including prevalence, load and virulence. | We combined the two previous versions of the study inclusion criteria to include several different measures of parasite success, which were later used for contextual factor analysis. |
| 2) Parasite success data was collected from two or more host populations with any comparable difference in genetic diversity, such as the level of relatedness among individuals (inbred versus outbred), genotypic diversity (high versus low) or heterozygosity. | - Ekroth, Rafaluk-Mohr and King, 2019: Data was collected from any study with two distinct populations and any measured difference in diversity.  - Gibson and Nguyen, 2020: Collected data for two or more populations. | We collected data from studies of multiple populations with any comparable difference in genetic diversity to increase our sample size and because there was one study with differences in genetic diversity which were not comparable between all pairwise combinations (Baer 2001). |
| 3) Genetic diversity was measured at the host population level and not community diversity or individual-level genetic heterozygosity. | - Ekroth, Rafaluk-Mohr and King, 2019: Used the exact same wording.  - Gibson and Nguyen, 2020: Stated that host genetic diversity had to be intra-specific. | We followed both Ekroth, Rafaluk-Mohr and King, 2019 and Gibson and Nguyen 2020 in this criterion. |
| 4) The study focused on an animal (or bacterial) host species. | - Ekroth, Rafaluk-Mohr and King, 2019: Excluded studies of agricultural systems.  - Gibson and Nguyen, 2020: Did not specify the study system. | We did not include any studies of non-animal populations, except for prokaryotic bacteria, because a more detailed analysis of the plant literature would require a separate review. |
| 5) The study does not re-analyze the data from a previously published study. | - Both Ekroth, Rafaluk-Mohr and King, 2019 and Gibson and Nguyen, 2020: Did not include this specification. | We included this specification because Ekroth, Rafaluk-Mohr and King, 2019 included data from two different studies by Baer and Schmid-Hempel which were based on the same dataset. |
| 6) The parasite success data was not replicated simply by using an alternate way of measuring host population diversity. | - Both Ekroth, Rafaluk-Mohr and King, 2019 and Gibson and Nguyen, 2020: Did not include this specification. | We included this specification because there two studies included by the previous meta-analyses (Giese 2003 and Puurtinen 2004) which included parasite success data for the same populations with two different measures of genetic diversity, which was a form of pseudoreplication. |
| 7) An attempt to take the parasite success data from clearly illegible figures was not made. | - Both Ekroth, Rafaluk-Mohr and King, 2019 and Gibson and Nguyen, 2020: Did not include this specification. | We included this specification because Gibson and Nguyen, 2020 had collected data from two studies with illegible figures (Agha, 2018 and van Houte et al. 2016). |
